# A low-autofluorescence, transparent resin for multiphoton 3D printing

**DOI:** 10.1101/2020.12.15.422922

**Authors:** George Flamourakis, Antonis Kordas, Georgios D. Barmparis, Anthi Ranella, Maria Farsari

## Abstract

Multiphoton lithography allows the high resolution, free-form 3D printing of structures such as micro-optical elements and 3D scaffolds for Tissue Engineering. A major obstacle in its application in these fields is material and structure autofluorescence. Existing photoresists promise near zero fluorescent in expense of poor mechanical properties, and low printing efficiency. Sudan Black B is a molecular quencher used as a dye for biological studies and as means of decreasing the autofluorescence of polymers. In our study we report the use of Sudan Black B as both a photoinitiator and as a post-fabrication treatment step, using the zirconium silicate SZ2080™ for the development of a non-fluorescent composite. We use this material for the 3D printing of micro-optical elements, and meso-scale scaffolds for Mesenchymal Stem Cell cultures. Our results show the hybrid, made photosensitive with Sudan Black B, can be used for the fabrication of high resolution, highly transparent, autofluorescence-free microstructures.

## Introduction

Multiphoton Lithography is an ultrafast laser-based additive manufacturing technique, which allows the fabrication of free-form 3D structures with submicron resolution [1]. Since firstly employed by the Kawata group in 1997 [2], MPL has been adopted as a powerful tool by a variety of research fields, ranging from optics and photonics [3], to biomedical and mechanical devices [4].

In general, the materials employed in MPL consist of a mixture of monomers and oligomers, that crosslink employing radical photopolymerization [5]. These radicals are provided by the photoinitiator (PI), a small molecule which is transparent at the laser wavelength *λ* (usually around 800 nm), absorbs at the two-photon wavelength *λ/2*(400 nm), has high two-photon cross-section, high radical quantum yield and generates highly-active radical species. Some of the PIs routinely used in MPL are Irgacure 369 [6], (4,4’-bis (dimethylamino) benzophenone) [7], and thioxanthone derivatives [8]. All these materials exhibit autofluorescence, even after the photopolymerization and subsequent development in a solvent [9].

This has hindered the application of the technology in research fields that employ fluorescence microscopy, a powerful and valuable tool in the hands of researchers, especially in biomedicine. Specifically, the multiple techniques of fluorescence imaging allows researchers to visualize the colorless, transparent tissues, cells, individual organelles and macromolecules, to study living cells and their intracellular communications and also to measure the pH, free calcium and NAD(P)H concentration in the cytoplasm [10]. It is therefore understandable that in the field of Tissue Engineering and Regenerative Medicine the development of scaffolds that will not impede or hinder the imaging of the generated tissues and organs is a necessity.

Optogenetics is an emerging, interdisciplinary research area which combines genetic and optical technologies to steer and monitor specific biological processes such as enzyme activities, protein conformation, membrane voltage, ions and molecules [11], [12]. Towards this scope, light-activated proteins, (optogenetic actuators or fluorescent sensor proteins) are genetically targeted to the cells of interest [13]. It is therefore apparent that the development of transparent non-fluorescent high-resolution 3D substrates is also applicable to optogenetics.

To address the drawback of autofluorescence alternative approaches have been used. To this end, there has been active research in producing low autofluorescence photoresists scaffolds and micro-optical elements for MPL, such as the IP-Visio resin by the leader in the field, Nanoscribe GmbH [14]. Other methods reported are: the photobleaching of polymers (a transient solution, as autofluorescence recovers after some hours of treatment) [15]; image processing, which involves digitally reducing the fluorescence background, reducing at the same time the signal emitted by the sample [16]; and also the use of infrared fluorophores which isn’t convenient as few microscopes are equipped with the appropriate filter and detector. It is therefore evident that a non-fluorescent free-standing scaffold is crucial for biomedical applications, as fluorescence labeling is used as a primary approach to investigate the interactions between cells and polymeric scaffolds at the molecular level.

Sudan Black B (SBB) [17] is a azobenzene-based dye widely used for the staining of lipids such as phospholipids, sterols and neutral triglycerides [18]. It has been shown that SBB can be used as an MPL photoinitiator [19], and that treating polymeric films with it improves fluorescence imaging of biological samples, by quenching natural autofluorescence [17], [20], [21], without affecting cell behavior. In this manuscript, we present our research into the suitability of SBB as a PI for the fabrication of low-autofluorescence cell scaffolds by MPL using lasers operating in the 800 nm region, where SBB absorption is negligible. We show that SBB can be employed for the fabrication of both high-resolution, complicated nano-structures and large-scale tissue engineering scaffolds using a zirconium silicate as an organic-inorganic resin. We seed the fabricated 3D scaffolds with Mesenchymal Stem Cells (MSCs), stain them with fluorescent dyes for the most common 3 colors (Red, Green, Blue) and show that the autofluorescence of the resulting structures does not interfere with the cell derived dye fluorescence, thus allowing detection of minimum signal intensities compared to other resists, when subjected to a further SBB treatment as described in [21]. Finally, we show that, even though the prepolymer is very dark, the polymerized thin films and 3D printed micro-optics are transparent, making this material suitable for the fabrication of 3D printed, autofluorescence-free optical components such as diffractive and micro-optical elements.

## Results

### Fluorescence of SBB-photosensitized scaffolds

The material employed for MPL fabrication is the organic-inorganic hybrid material SZ2080™, reported previously [22]. Two photoinitiators were used in this study: SBB, for the fabrication of low-autofluorescence 3D structures, and 4,4’-bis(diethylamino) benzophenone (Michler’s ketone), as a control. For brevity, the SBB-photosensitized composite will be called SBB-C, while the Michler’s ketone-photosensitised composite will be called BIS-C.

To test the overall fluorescence of the new material SBB-C, a series of 3D woodpile scaffolds were fabricated employed both SBB-C and BIS-C as a control. These scaffolds were designed using the CAD software Fusion 360 (Autodesk, California, USA) to have dimensions 320μm × 264μm × 80μm.

We obtained fluorescence and SEM images of scaffolds fabricated with BIS-C and SBB-C before and after SBB treatment, at the Red, Green, and Blue channels. The results are presented in **Figure 1**.

**Figure 1.**
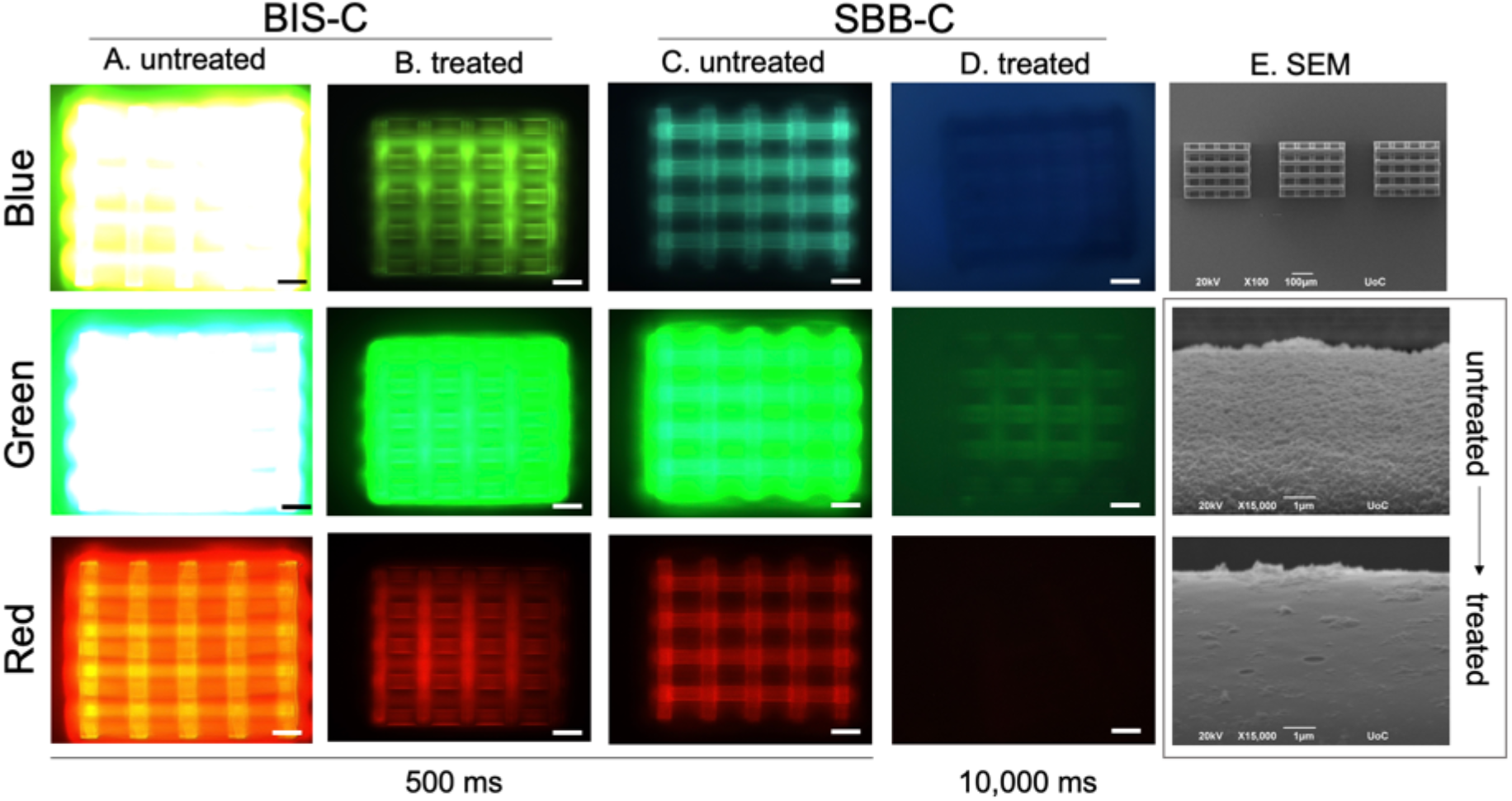
Comparative study of the fluorescence of BIS-C and SBB-C. Untreated scaffolds demonstrated the highest fluorescence, while treated SBB samples exhibited minimum fluorescence. The last column depicts SEM images of the scaffolds in top. Further magnification 15,000X reveals the effect of treatment on the polymer. Scale bar corresponds to 50μm.

Column (a) shows fluorescent images of scaffolds made using the BIS-C hybrid. Scaffold autofluorescence was so high that the green channel penetrated the blue one, making the scaffold glow green in both channels (exposure: 500ms). Even after treatment with SBB (column (b)), autofluorescence is very high, for the same exposure time. The SBB-C hybrid exhibits much lower autofluorescence (column (c)); any residual autofluorescence is most likely due to ZrO_2_ nanoparticles formed in the resin during photopolymerization. In column (d) we show scaffolds fabricated using the SBB-C hybrid, and subsequently treated with SBB. Note that the camera exposure is 10,000ms, as a 500ms exposure was not sufficiently long for the scaffolds to be visible. Even at this longer exposure, autofluorescence levels are negligible and the scaffolds were barely visible, highlighting the efficiency of the SBB-C/SBB treatment combination.

Column (e) shows SEM images of the SBB-C scaffolds and their surface before and after SBB treatment. It is clear that the SBB treatment does not damage or in any way affect the integrity of the scaffolds. Furthermore, the high magnification SEM image (15,000x) shows that the SBB treatment leads to much smoother surfaces, where SBB has been adsorbed inside the polymer pores, as proposed by Jafaar *et al* [17].

This effect was further investigated using the ImageJ software where the mean fluorescence intensities were measured (**Figure 2**). Green and blue color of BIS-C, untreated samples demonstrated the highest intensities of 250au while the red color showed a much lower intensity at around 150au. Treatment with SBB lowered the intensities of the BIS-C samples to comparable levels to the SBB-C untreated ones. Further treatment of SBB-C samples eliminated fluorescence almost completely. It is remarkable that almost 90% autofluorescence reduction between BIS-C untreated and SBB-C treated samples was observed.

**Figure 2.**
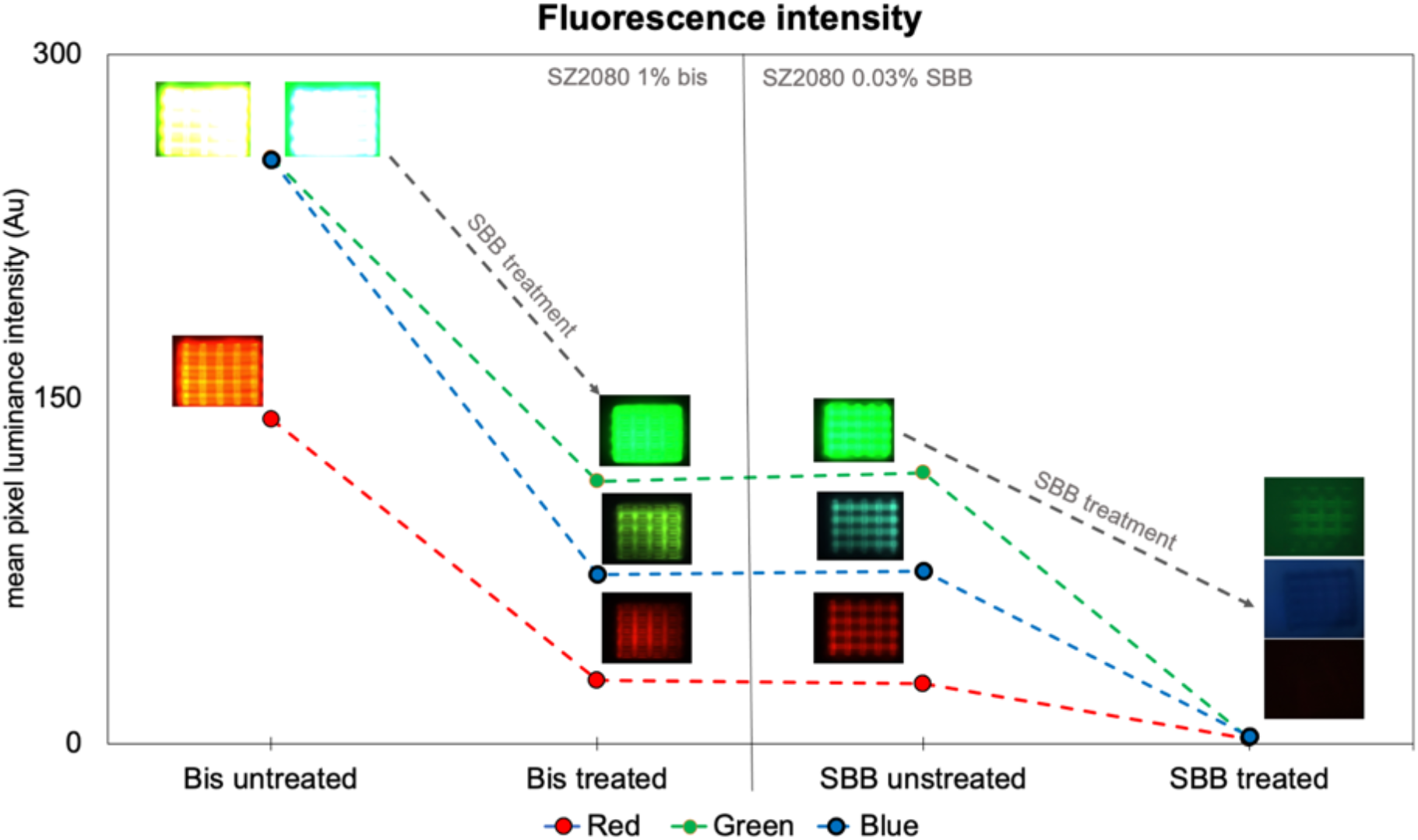
Fluorescent intensities of BIS-C and SBB-C samples. Exposure time was set at 500ms with the exception of SBB treated samples, where exposure time was set at 10,000ms due to the inability to observe the scaffolds at lower exposure times. Remarkably, Bis treated and SBB untreated samples exhibit comparable intensities, whereas the intensities of SBB treated samples at 500ms would be almost zero.

### Cell studies

As discussed earlier, the low autofluorescence of the SBB-C hybrid makes it an ideal material candidate for 3D scaffold fabrication, for the investigation of 3D cell cultures using fluorescence techniques. To this end, the woodpile geometry was chosen as our initial scaffold fabrication, as it provides both complexity and large surface areas for cells to grow on. After 3 or 4 days of culture, MSCs reached confluence of 90% and for their staining were used three different dyes that fluoresce at different wavelength. More specifically, we used DAPI dye fluoresce at 358nm for the nucleus staining and the FITC conjugated phalloidin 568 as a high-affinity filamentous actin (F-actin) probe. Finally, the secondary anti-rabbit green (488 nm) conjugated antibody utilized to detect the primary YAP antibody (YAP) that was previously bound to the YAP protein isoforms. Each one corresponds to blue, red and green channels respectively. There were 4 different scaffolds prepared in total; woodpiles fabricated with BISC both treated and untreated with 0.3% SBB, and woodpiles made with SBB-C both treated and untreated with 0.3% SBB. It should be noted that the treatment took place right before cell seeding for 1h in Room temperature (RT).

**Figure 3** clearly demonstrates the suitability of the SBB-C hybrid for scaffold fabrication regarding cell growth for various types of 2D and 3D studies. Cells show good survival rates, comparable with already established biomaterials such as gelatin methacryloyl (GELMA) hydrogels [23], poly(lactide-co-glycolide) (PLGA), polycaprolactone (PLA) etc. [24], this is backed up by the live/dead assay results (Figure 3. I,J). It can be seen that cell viability reached 99%. This is not surprising, as both the zirconium silicate and SBB are known to show good compatibility with cells [25].

**Figure 3.**
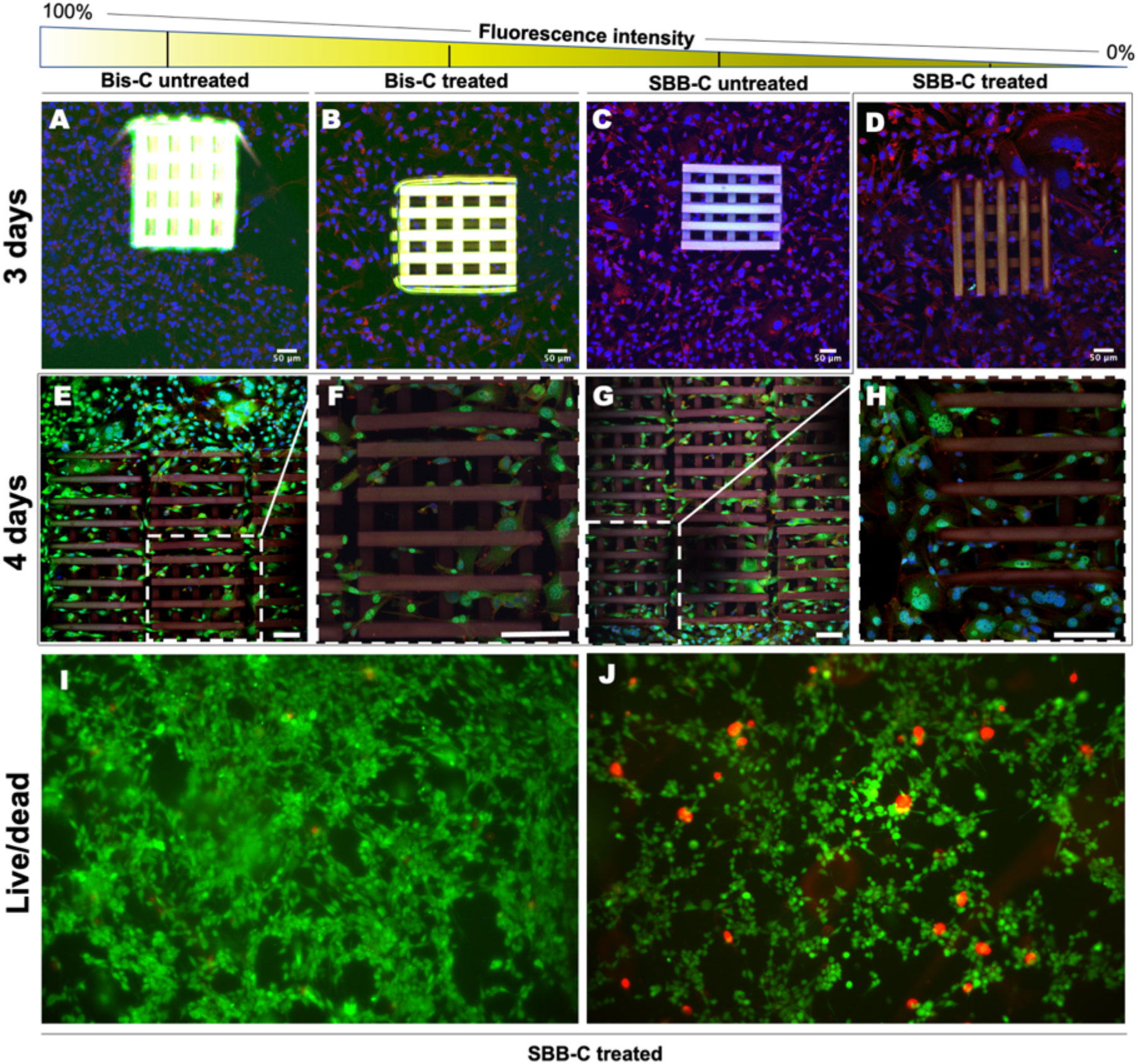
Confocal Microscopy images of the woodpile scaffolds seeded with MSCs (**A-H**). Each image is a composite of all z-stacks and merged channels. Images of live/dead experiment of MSCs on thin SBB-C films in 48h time point (**I, J**). **A**) BIS-C, untreated; highest fluorescence. Evaluation of the cell population inside was impossible. **B**) BIS-S, treated; significantly lower fluorescence resulting in some cells to be visible. **C**) SBB-C, untreated; low fluorescence as many cells are clearly visible inside the woodpile. **D**) SBB-C, treated; lowest fluorescence. The nuclei and the actin of the cells inside the scaffold are clearly visible. Cells that adhere on the top of the scaffold are distinguishable from those inside and under it. **E**) Image of 9 woodpiles with MSCs with the respective **F**) zoomed image of the middle scaffold. **G**) Upper end of the same 9 woodpile scaffolds with **H**) zoomed image of the left-end woodpile. **I, J**) 48h of MSCs culture on SBB-C film. Green represent live cells and red the dead ones.

Using the SBB-C for scaffold fabrication, and subsequently treating these scaffolds with SBB allowed us to eliminate scaffold autofluorescence. During confocal microscopy imaging, cells can be clearly distinguished both inside the pores of the scaffold and on its surface. The significance of these results is profound, as scaffold autofluorescence has been identified as a major hurdle in employing confocal microscopy in 3D cell studies [26]. Natural cell environments are three-dimensional and information gained from 2D or 2.5D studies may only be partial or incomplete. By allowing 3D imaging, the user can have a global understanding on cell behavior and interactions of cells with the scaffolds. The incorporation of confocal microscopy for imaging can be crucial and beneficial for fields such as tissue engineering when compared to alternative imaging techniques like SEM, which have no coloring options during observation, are not suitable for live imaging, and provide no information on the cell interior on molecular level.

Even though the SBB treatment reduced autofluorescence dramatically, is still noticeable. We believe this is due to the scaffold attachment to the substrate; the interface between the glass substrate and the scaffold is not reachable for SBB treatment thus allowing some fluorescence to be detected.

To investigate the suitability of SBB-C for the fabrication of high resolution, complex geometries, we employed two very demanding metamaterial designs found in literatures: the negative Poisson’s ratio auxetic scaffold honeycomb reentrant (or bowtie scaffold) [27] and the positive Poisson’s ratio ultra-stiff, ultra-light structure tetrakaidecahedron (often refer as Kelvin foam) [28]. Those geometries were fabricated using the ‘Nanocube’ setup in a unit-cell to unit-cell manner avoiding slicing and ensuring their best mechanical stability and resolution. The auxetic unit cell pore size was 50μm while the Kelvin cell pore size was 95μm, allowing us to create two different conditions for the cell culture, one 2.5D auxetic scaffold (total dimensions of 2mm × 2mm × 100 μm) and a 3D Kelvin foam where the cells could penetrate the unit cells (total dimensions of 4mm × 4mm × 95 μm). The large dimensions of both scaffolds ensured that a good number of cells would be seeded on them creating the perfect mechanical micro-environment for our culture. After 3 days of culture, both samples were stained and prepared for confocal observation as before. This time the observation was conducted from top to bottom in order to avoid the untreated interface regions of the scaffolds.

For the first time it was possible to observe stained MSCs inside such complicated architectures and quantify their interactions with the scaffolds at molecular level by visualizing the levels of YAP protein in the green channel (**Figure 4**). The whole 2mm × 2mm × 100μm scaffold was mapped but only a portion of it is shown for convenience. MSCs can be spotted all around both scaffolds and by using higher magnification, it is clear that cells bend and reform the auxetic scaffold in agreement with results acquired in previous work [7], indicating similar mechanical response to those of BIS-C. The Kelvin foam scaffold is a very stiff geometry; thus, no deformations were observed. It is remarkable how further magnification using a 63x lens can make the scaffold barely visible, making it possible to focus the observation solely on the cells’ responses to it without the limitations of background fluorescence in all 3 different colors. The unit cell is large enough for many cells to enter as it can be seen in the zoomed unit cell. Furthermore, multiple phenotypes of MSCs are distinguishable on the same scaffold where YAP protein is located mostly in the nucleus (hinting that the extracellular mechanical signal was transferred to the nucleus, inducing gene transcription) or in the cytoplasm with incomparable detail [29]. Alongside with the treated meta-material scaffolds, untreated ones were fabricated and prepared for SEM observation with exactly the same culture conditions as the confocal ones. Those images acquired from SEM were used as reference.

**Figure 4.**
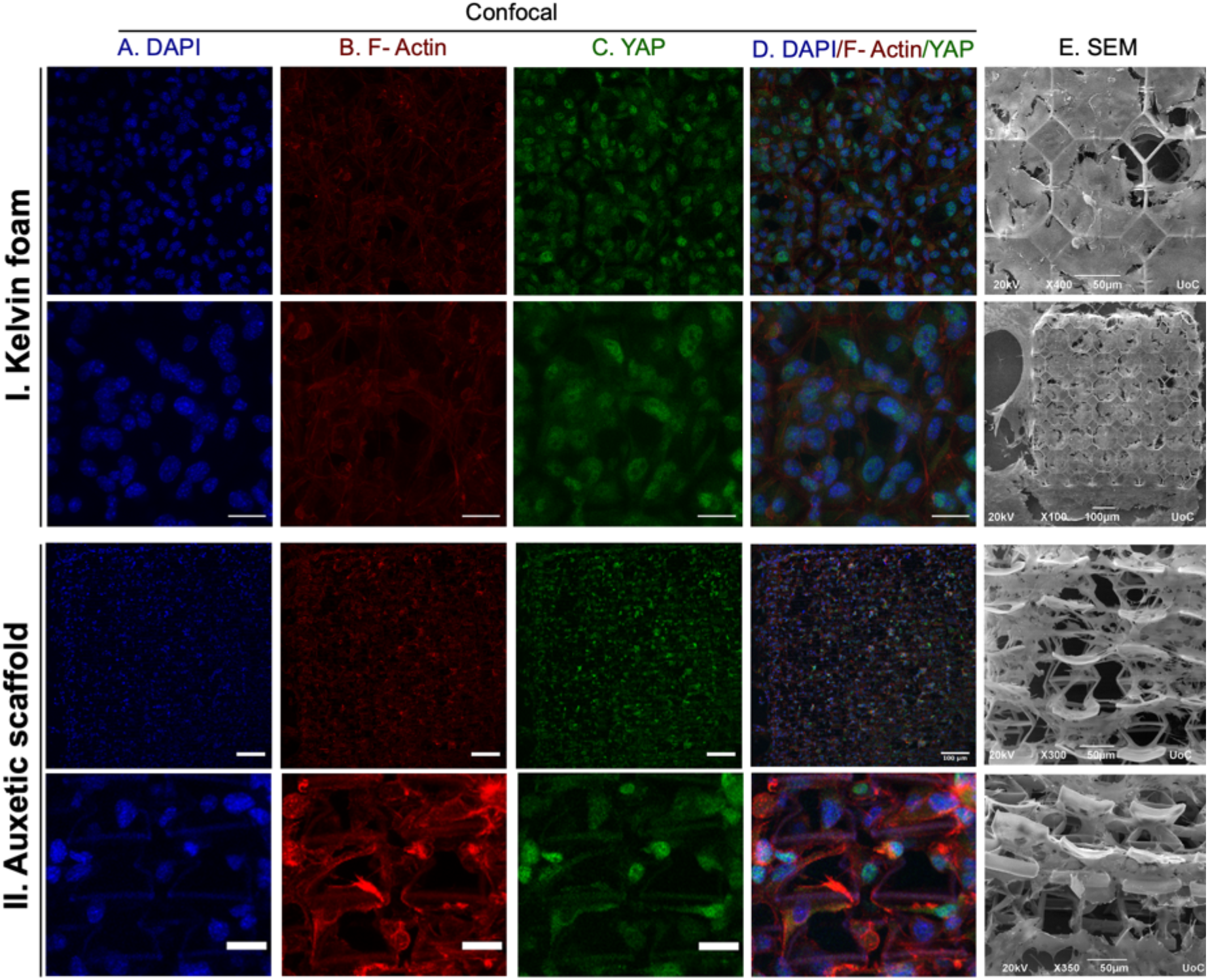
MSCs culture in auxetic scaffold and Kelvin foam made with SBB-C treated material under confocal microscopy and SEM. The first row shows a general view of the kelvin foam with scale bar corresponding to 50μm. The second row shows a zoomed area of a single unit cell with several cells inside (scale bar 20μm). Third row shows a general view of the Auxetic scaffold (scale bar 100μm) and forth row a zoomed picture of individual unit cells (scale bar 20μm). **(A) Blue channel:** nuclei, DAPI. **(B) Red channel**: F-actin, phalloidin 568. **(C) Green channel:** YAP protein, FC488. **(D)** All 3 channels merged together. All images have z-projected all the stacks. **(E)** SEM images.

### Transparent Micro-optical elements

While the SBB-C hybrid is almost black in liquid form, after photopolymerization and development it becomes clear, as shown in **Figure 5**. The material transmission is approximately 90% over the visible part of the spectrum, comparable to commonly used materials for the fabrication of optical elements, such as ORMOCLEAR®[30]. This high transmission, combined with the SBB-C low autofluorescence and high structuring accuracy, makes it an ideal photopolymer for the fabrication of micro-optical elements (MOEs). To this end, two different 100micron-diameter MOEs were fabricated (inserts in **Figure 5**), a high NA lens (insert b) and a beam expander (insert c), as previously described by Dietrich *et al.* using the Nanoscribe photoresist IP-vision [31].

**Figure 5.**
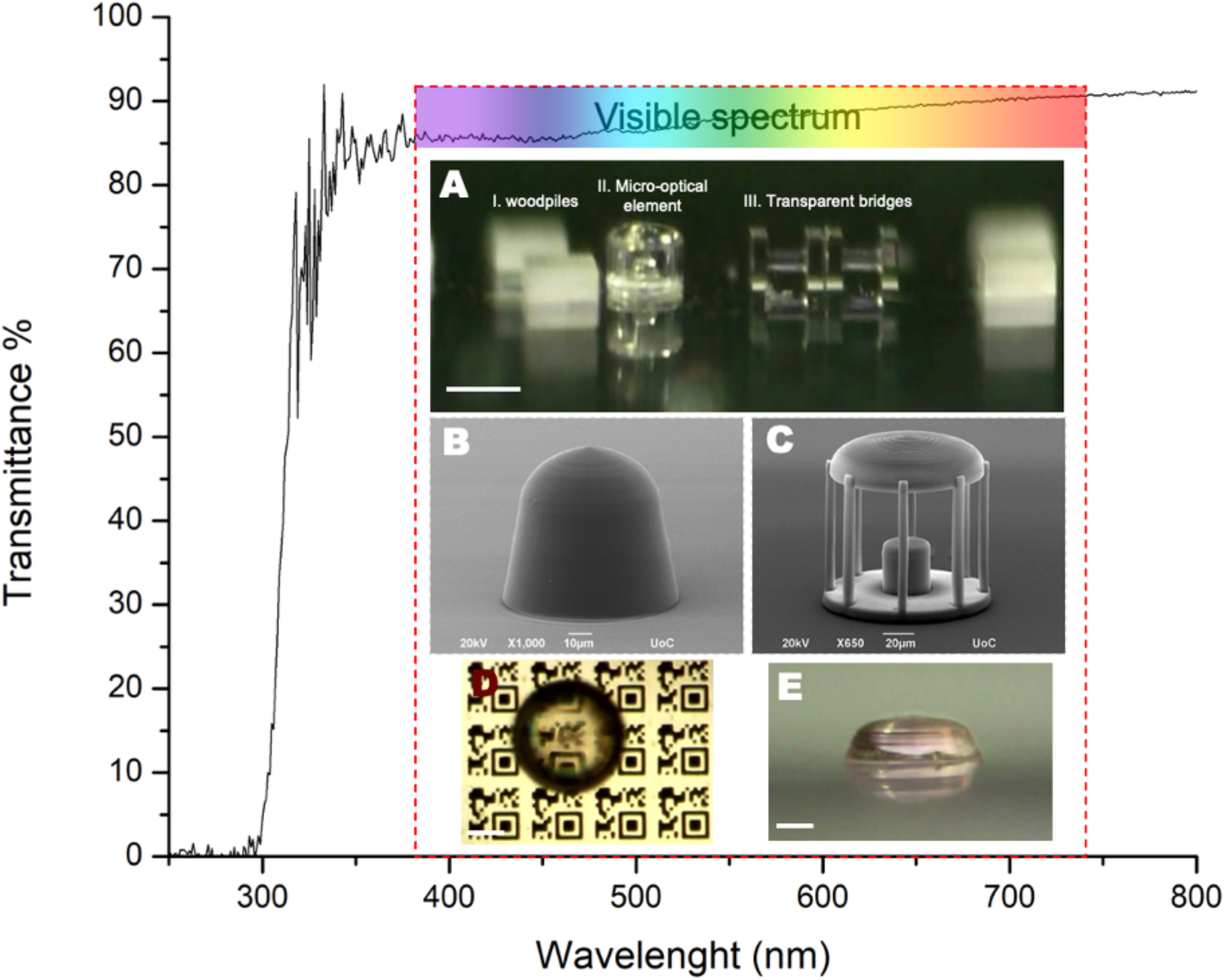
Transmittance percentage over wavelength of a thin, spin-coated SBB-C film. High transmittance is observed for the entirety of the visible spectrum that only drops to around 0% in the Ultra-violet region. A) Enclosed are the images of various structures fabricated with SBB-C where absolute transparent was achieved. B, C). SEM images of the 2 micro-optical elements. D) High NA optical lens placed over a printed QR code and its’ respective E) front view. Scale bars correspond to 100μm.

For the design of these elements the CAD software Fusion 360 was used, while the fabrication was carried out using the ‘Galvo’ set-up. In this case, an oil-immersed microscope objective lens (40x, NA=1.4, Zeiss, Plan Apochromat) was employed, and the printing conditions were: laser output = 30mW (measured before the objective); galvo scanning speed= 90mm.s^−1^; hatching distance= 0.2μm and slicing distance = 0.2μm. SEM images show that the surfaces of the optics are smooth, and the structures have been printed with accuracy, demonstrating the suitability of SBB-C for optics fabrication. Furthermore, we acquired front facing pictures using a Dino-lite digital microscope at 220x magnification (AM7915MZT, AnMo Electronics Corporation, Taiwan) to investigate the transparency of the optics. Insert A shows 3 different structures, (I), small woodpile structure, (II) the beam expander micro-optical element and (III), 2 bridges fabricated for mechanical investigation purposes. It seems that all structures all completely transparent and almost glass-like, fact that is further investigated in insert D, where the high NA lens was placed right above a printed QR code, making it totally visible.

## Discussion

In the present study, SBB was used as a photoinitiator in low concentration (0.03% w/v) with the well-established material SZ2080 and as a post-fabrication treatment process to reduce and even eliminate the autofluorescence of polymer scaffolds. We were able to print very sophisticated and demanding 3D designs for both biomedical and micro-optical applications. Those designs included bulk, large woodpile structures made using galvanometric mirror system in very high scanning speed, meso-scale complicated mechanical metamaterial scaffolds fabricated in a unit cell to unit cell manner employing piezoelectric stages and very high resolution micro-optical elements. In addition to excellent resolution in combination with ultra-fast scanning speeds, we showed that the SBB-photosensitized hybrid is suitable for printing low autofluorescence 3D structures, which becomes negligible when combined with further treatment of the structures with SBB. Using confocal microscopy for low auto fluorescent materials has always been a challenge for fields like tissue engineering, since most of the commercially available materials are limited to 2D or 2.5D scaffolds, limiting their use in 3D applications; this, however, does not seem to be a limiting factor here, as fabrication of large, complicated 3D scaffolds was possible for MSCs cultures. Moreover, the seeded MSCs showed excellent survival rates as no cytotoxicity was observed *(**fig. 2, I, J***). Furthermore, the use of SBB-C was expanded to the fabrication of two micro-optical elements, a high NA lens and a beam expander; both structures maintained high resolution after fabrication, which combined with the high transmission of SBB-C in the visible spectrum and low autofluorescence, highlights the potential of our approach in optics.

Collectively, the results demonstrate the suitability of the SBB-C hybrid material for MPL by improving the fluorescence of the cell-polymer scaffolds complexes allowing high decree observation of the dynamic interplay between the extracellular environment and the cells in molecular level and we expect that SBB-C should be suitable for a plethora of applications outside those described here.

## Methods

### Photoresists preparation

The material employed for MPL fabrication is the organic-inorganic hybrid material SZ2080™, reported previously [22]. In brief, methacryloxypropyl trimethoxysilane (MAPTMS, 99%, Polysciences Inc.) was hydrolyzed with dilute HCl. Separately, methacrylic acid (MAA, 98%, Sigma-Aldrich) and zirconium n-propoxide (ZPO, 70% in propanol, Sigma-Aldrich) were combined in a molar ratio of 1:1 and stirred for 30 min; then the second composite was slowly added to the first. Two photoinitiators were used in this study: SBB (Sigma-Aldrich 0.03% w/v), for the fabrication of low-autofluorescence 3D structures, and 4,4’- bis(diethylamino) benzophenone (Michler’s ketone, Sigma-Aldrich, 1% w/w), as a control. It should be noted that Michler’s ketone was directly incorporated inside the material in 1% wt ratio, while SBB was firstly mixed with isopropanol in a concentration of 0.3% w/v, and then through a 1:10 dilution was added to the material (final concentration of 0.03% wt). After adding the photoinitiator, all materials were filtered using a 0.2μm pore filter.

### Sample preparation

3D structures were fabricated using MPL, while thin films were fabricated via spincoating. Prior to structure fabrication, the glass substrates were lyophilized to improve material adhesion. In brief, 13mm glass coverslips were placed in a glass container filled with 93%ethanol, and sonicated for 1h. They were subsequently immersed in a mixture of 1:80 dichloromethane: MAPTMS, and were sonicated for a further 4 hours. Next, they were again immersed in 70% ethanol, for another hour of sonication. They were stored in fresh ethanol, and dried before use. The procedure was carried out at Room Temperature (RT).

For thin film preparation, 15μL of resist were deposited in the center of a coverslip and the material was spin-coated at 4000rpm for 40sec with an acceleration step of 500rpm/sec. The films were then either heated at 80°C for 1h or placed in vacuum for 48h, and were left overnight under a UV lamp. Afterwards, they were immersed in 1:1 4-methyl-2- pentanone/isopropanol, washed in isopropanol/dd-H_2_O for 2 min and left to dry.

For the 3D structure fabrication, 15μL of resist were deposited in the center of each coverslip, and the samples were heated at 95°C for 1h, for the composite to condense and become a hard gel. The 3D structures were subsequently fabricated using MPL. After fabrication, the samples were developed in 4-methyl-2-pentanone/Isopropanol (1:1) for 5min and then washed in isopropanol/dd-H_2_O for 2 min to remove any unpolymerized material and were left to dry.

### Structure Fabrication

The MPL experimental setup employed in these experiments consists of two scanning systems, which from now on will be referred to as the ‘Nanocube’ system and the ‘Galvo’ system. Both have been described in detail in [32]. The main difference between them is that in the ‘Nanocube’ system the laser beam is fixed in a centric position, and 3D writing is achieved by moving the sample using piezoelectric and linear stages (PI), while in the ‘Galvo’ set-up, the beam moves in the xy plane by galvanometric mirrors, and linear stages are used for z-axis and large-scale movement. Both systems use the same irradiation source, a femtosecond fiber laser (FemtoFiber pro NIR, Toptica Photonics AG). Woodpile structures were made using the ‘Galvo’ system, while auxetic and tetrakaidecahedron (Kelvin cell) structures were made using the ‘Nanocube’ system. the optimized printing conditions of each structure are summarized in ***Table 1***.

**Table 1.** Printing conditions of the various structures.

| Structure | Objective lens | System | Scanning speed ($\mu\text{m.s}^{-1}$) | Hatching distance ($\mu\text{m}$) | Slicing distance ($\mu\text{m}$) | Energy output (mW) |
| --- | --- | --- | --- | --- | --- | --- |
| Woodpiles | 20x | GALVO | 50,000 | 0.5 | 5 | 20 |
| Kelvin foam | 20x | NANO | 100 | - | - | 30 |
| Auxetic scaffold | 40x | NANO | 100 | - | - | 25 |
| Optical elements | 40x oil | GALVO | 90,000 | 0.2 | 0.2 | 30 |
| Bridges | 40x oil | GALVO | 90,000 | 0.2 | 1 | 20 |

### SBB treatment

To eliminate any remaining autofluorescence, some of the 3D woodpile structures and thin films were treated with SBB following the procedure described in [21]. In brief, the samples were immersed in a solution of 0.3% SBB in 70% EtOH for 1h followed by several washings with EtOH.

### Cell culture

Mouse Bone Marrow Mesenchymal Stem Cells (BM-MSCs, Cyagen, California, USA) were used for all experiments as they are widely considered the golden standard in tissue engineering and regenerative medicine. The cells were incubated at 37°C in an atmosphere of 5% CO_2_ in air in Low-glucose Dulbecco’s Modified Eagle’s Medium (DMEM, Gibco, Invitrogen, Karlsruhe, Germany) supplemented with 10% heat inactivated Fetal Bovine Serum (FBS, Gibco, Invitrogen, Karlsruhe, Germany) and 1% penicillin/streptomycin (Gibco, Invitrogen, Karlsruhe, Germany). The 3D scaffolds were sterilized by immersing the coverslips in 70% ethanol and then drying under UV light for 30min. The sterilized scaffolds were placed in a 24-well plate (Sarstedt, Numbrecht, Germany), seeded with 5.10^4^ cells/ml in a total volume of 1ml (p.3-4) and incubated at 37°C for 3 days in the same medium described above. Seeding took place after a confluent flask of 90%.

### Immunofluorescence Staining

Upon the end of cultivation time (both 3 and 4 days) the medium was removed and each well was washed three times with 1x Phosphate Buffered Saline (PBS, Gibco, Invitrogen, California, USA) followed by fixation with 4% w/v Paraformaldehyde (PFA, Sigma Aldrich, Missouri, USA in PBS for 15 min. The cells were then permeabilized with Triton X-100 0.5% v/v (Sigma Aldrich, Missouri, USA) in PBS for 15 min, followed by blocking with 2% w/v Bovine Serum Albumin (BSA, Biofroxx, Einhausen, Germany) in PBS for 1h. Rabbit YAP antibody (Yes Associated Protein, YAP65, 1:100 dilution, Cell Signaling Technology®, Danvers, USA) was used to dye all the isoforms of YAP protein in 0.5% w/v BSA, 0.1% v/v Triton X-100 in PBS at 4°C overnight. The next day, TRITC-conjugated phalloidin-568 (EMD Millipore, Burlington, USA) (1:1,000 dilution) was used to dye the actin cytoskeleton alongside with CF488 anti-rabbit IgG antibody (Biotium, Fremont, USA, 1:500 dilution) in 0.5% w/v BSA 0.1% v/v Triton X-100 in PBS for 1h. Finally, the coverslips were placed upside down on a glass slide with slow fade mounting medium containing 4’,6-diamidino-2- phenylindole (DAPI, Life Technologies, Carlsbad, CA, USA) and kept at 4 °C until observation. It should be noted that between each step the samples were washed thrice with 1x PBS. All steps of the procedure were carried out at RT, unless otherwise stated. An inverted Confocal Microscope (Leica SP8 inverted confocal, Leica Microsystems) was used for observation of all samples.

### Scanning Electron Microscopy

To prepare samples for scanning electron microscopy (SEM) the cells after the desired timepoints were fixed using the cell fixation/preparation for SEM analysis protocol. Briefly, the culture medium was removed and the cover slips were washed twice with 1x PBS. Then, 2.5% v/v glutaraldehyde (GDA, Sigma Aldrich, Missouri, USA)/ 2.5% v/v paraformaldehyde (PFA, SigmaAldrich, Missouri, USA) in 1x PBS, incubated for 15min in room temperature (RT) followed by two washes with PBS 1x. Finally, the specimen was dehydrated in graded ethanol solutions (from 30% to 100%), underwent a critical point drying process (Baltec CPD 030) and 10 nm gold-sputter coat (Baltec CPD 050). Micro optical elements were only gold-sputtered post-fabrication with 10nm gold and observed.

### Live/dead cell assay

To investigate the effect of SBB as a PI and as a treatment (0.3% w/v in EtOH) on cell viability, a live/dead assay was performed on an MSC culture (p.3-4), seeded on SBB treated thin films of the SBB-C composite. A total number of 10.10^4^ cells were initially seeded on the films and cultured inside an incubator (5% CO_2_ at 37°C) for 2 days. Cell viability was examined by labeling live and dead cells using Live/Dead^®^ kits (Life Technologies, Carlsbad, CA, USA). Briefly, cells were incubated in PBS with ethidium homodimer-1 (EthD-1, 4 μM) and calcein AM (2 μM) for 40 min following the manufacturer’s instructions. Live cells were labeled with calcein AM (green) and dead cells were labeled with ethidium homodimer-1 (red) and observed under an epifluorescent microscope (Carl Zeiss, Axioscope 2 Plus).

## Acknowledgments

We acknowledge support by the projects “Advanced Research Activities in Biomedical and Agro alimentary Technologies” (MIS 5002469) which is implemented under the “Action for the Strategic Development on the Research and Technological Sector”, funded by the Operational Program “Competitiveness, Entrepreneurship and Innovation” (NSRF 2014-2020), HELLAS-CH (MIS 5002735) implemented under “Action for Strengthening Research and Innovation Infrastructures”, funded by the Operational Program “Competitiveness, Entrepreneurship and Innovation” (NSRF 2014-2020) both co-financed by Greece and the European Union (European Regional Development Fund), and PULSE, the European Union’s Horizon 2020 research and innovation program under grant agreement No 824996. The authors would also like to thank Dr. Phanee Manganas for her valuable help operating the confocal microscope, and Ms. Aleka Manousaki, for expert help with SEM.

## Authors contributions

G.F. conceived the project, designed, optimized and fabricated the scaffolds and performed the cell culture experiments using the mechanical metamaterial scaffolds alongside with the SEM experiments. A.K. performed the immunofluorescence staining and the rest cell culture experiments. Both G.F. and A.K. contributed to the acquisition and analysis of data. G.B. developed the 3D printing program used for the ‘Galvo’ setup. M.F. and A.R. designed the work and supervised the project. All authors contributed to the writing of the manuscript.

